# Epidermal Growth Factor Receptor Signaling Governs the Host Inflammatory Response to Invasive Aspergillosis

**DOI:** 10.1101/2024.09.10.612305

**Authors:** Hong Liu, Jianfeng Lin, Quynh T. Phan, Vincent M. Bruno, Scott G. Filler

## Abstract

The epidermal growth factor receptor (EGFR) has been identified as an epithelial cell receptor for Mucorales fungi and *Candida albicans*. Blocking EGFR with small molecule inhibitors reduces disease severity in mouse models of mucormycosis and oropharyngeal candidiasis. In contrast, cases of invasive aspergillosis have been reported in cancer patients who were treated with EGFR inhibitors, suggesting that EGFR signaling may play a protective role in the host defense against this infection. Here, we analyzed transcriptomic data from the lungs of mice with invasive aspergillosis and found evidence that *Aspergillus fumigatus* infection activates multiple genes that are predicted to function in the EGFR signaling pathway. We also found that *A. fumigatus* infection activates EGFR in both a human small airway epithelial (HSAE) cell line and in the lungs of immunosuppressed mice. EGFR signaling in HSAE cells is required for maximal endocytosis of *A. fumigatus* and for fungal-induced proinflammatory cytokine and chemokine production. In a corticosteroid immunosuppressed mouse model of invasive pulmonary aspergillosis, inhibition of EGFR with gefitinib decreased whole lung chemokine levels and reduced accumulation of phagocytes in the lung, leading to a decrease in fungal killing, an increase in pulmonary fungal burden, and accelerated mortality. Thus, EGFR signaling is required for pulmonary epithelial cells to orchestrate the host innate immune defense against invasive aspergillosis in immunosuppressed hosts.

**Importance:** When *A. fumigatus* infects the lungs, it invades epithelial cells that line the airways. During this process, the fungus interacts with epithelial cell receptors. This interaction stimulates epithelial cells to endocytose the fungus. It also induces these cells to secret proinflammatory cytokines and chemokines that recruit phagocytes to the site of infection where they can kill the fungus. Here, we show that in small airway epithelial cells, the epidermal growth factor receptor (EGFR) acts a sensor for *A. fumigatus* that triggers the production of chemokines in response to fungal infection. In corticosteroid-immunosuppressed mice, blocking EGFR with the kinase inhibitor, gefitinib reduces chemokine production in the lungs. This leads to decreased accumulation of neutrophils and dendritic cell in the lungs, reduced *A. fumigatus* killing, and increased mortality. These results provide a potential explanation as to why some cancer patients who are treated with EGFR inhibitors develop invasive aspergillosis.

## INTRODUCTION

Conidia of the fungus *Aspergillus fumigatus* are ubiquitous and inhaled by most individuals every day. After inhalation, the conidia are deposited in the distal airways and alveoli. In healthy individuals, the conidia are cleared by neutrophils and pulmonary macrophages. In patients who are immunocompromised due to the presence of neutropenia, administration of high-dose corticosteroids, or hematopoietic stem cell or organ transplantation, conidia can germinate into filamentous hyphae that cause invasive aspergillosis. Even with contemporary antifungal therapy, approximately thirty per cent of patients with invasive aspergillosis die (1, 2). Mortality associated with this disease is even higher when the fungus is resistant to mold-active triazole antifungal drugs, which constitute the mainstay of therapy (3).

Epithelial cells that line the airways and alveoli are among the first host cells to encounter *A. fumigatus.* Besides being targets of fungal adherence, invasion, and damage, these cells play a key role in orchestrating the host innate immune response to invasive aspergillosis (4–6). By secreting cytokines and chemokines, infected pulmonary epithelial cells recruit phagocytes to foci of infection and enhance their fungicidal activity, thus facilitating fungal clearance (7). Although it is known that pulmonary epithelial cells express pattern recognition receptors such as dectin-1 (5, 8), there is incomplete understanding of how these cells sense and respond to fungal pathogens such as *A. fumigatus*.

The epidermal growth factor receptor (EGFR) has been identified as an epithelial cell receptor for Mucorales fungi and *Candida albicans*. Blocking EGFR with small molecule inhibitors reduces disease severity in mouse models of mucormycosis and oropharyngeal candidiasis (9–11). In contrast, cases of invasive aspergillosis have been reported in patients with cancer who were treated with EGFR inhibitors including gefitinib, afatinib, erlotinib and osimertinib (12–15). These results suggest that EGFR may play a protective role in the host defense against invasive aspergillosis.

Here, we analyzed transcriptomic data from the lungs of mice with invasive aspergillosis and found evidence that *A. fumigatus* infection activates the EGFR signaling pathway. We also found that *A. fumigatus* infection stimulates EGFR in both a human small airway epithelial (HSAE) cell line and immunosuppressed mice with invasive pulmonary aspergillosis. EGFR signaling in HSAE cells is required for maximal endocytosis of *A. fumigatus* and fungal-induced secretion of proinflammatory chemokines and cytokines. In immunosuppressed mice with invasive pulmonary aspergillosis, inhibition of EGFR with gefitinib decreased whole lung chemokine production and reduced accumulation of phagocytes, which was associated with less fungal killing, impaired survival and increased pulmonary fungal burden. Thus, EGFR signaling is required for the host defense against invasive aspergillosis in the immunosuppressed host.

## RESULTS

### *A. fumigatus* infection activates EGFR in mice

To gain a comprehensive view of the host response during invasive aspergillosis, we performed RNA-seq analysis of the lungs of mice that had been immunosuppressed with cortisone acetate and then infected with *A. fumigatus* strain Af293 for 2, 4, and 6 days (16). Previously, we demonstrated that infection-induced changes in the host transcriptome can be used to identify signaling pathways that govern the interaction between the host and a fungal pathogen (17–19). Using the Ingenuity Pathway Analysis software (Ingenuity systems; http://www.ingenuity.com), we performed an Upstream Regulator Analysis on the sets of host genes that were differentially expressed (log_2_ fold-change >1 <-1, false discovery rate < 0.05, Tables S1-4) in the infected mice and the appropriate time-matched negative control groups, which were immunosuppressed but not infected.

The analysis indicated that the breadth of the host transcriptional response to *A. fumigatus* infection increased progressively over time (Fig. 1A). Multiple pathways known to be required for the host defense against invasive aspergillosis were predicted to upregulated in the infected mice by day 6 post-infection. These pathways included TNF-α, CSF2 (GM-CSF), IL-1α, IL-1β, STAT3, IFN-γ, and IL-17A (20–25).

**FIG 1.**
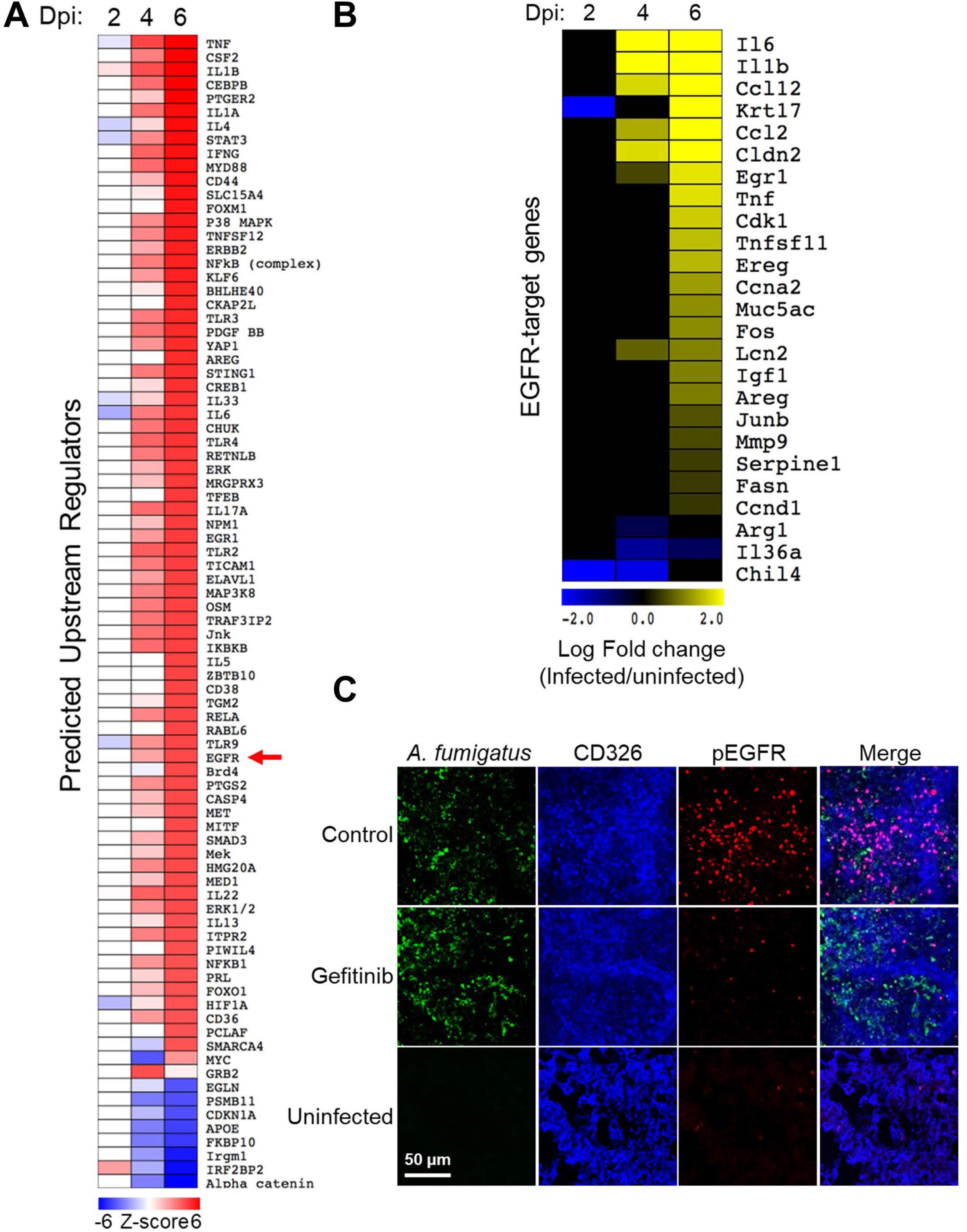
*A. fumigatus* infection of immunosuppressed mice activates the epidermal growth factor receptor (EGFR) in the lungs. (**A**) Heatmap showing the results of upstream regulator analysis of the transcriptome of the lungs of mice that had been immunosuppressed with cortisone acetate and infected with *A. fumigatus* for 2, 4, and 6 days. Control mice were immunosuppressed but not infected. DPI, days post-infection (**B**) Heatmap showing the effects of *A. fumigatus* infection on the mRNA levels of 25 genes in the EGFR pathway (**C**) *A. fumigatus* infection of immunosuppressed mice stimulates EGFR autophosphorylation, which is reduced in mice treated with the EGFR inhibitor, gefitinib. Confocal microscopic images of thin sections of the lungs of immunosuppressed mice 12 h after intratracheal inoculation with *A. fumigatus.* The sections were stained for *A. fumigatus* (green), pulmonary epithelial cells (CD326; blue), and phosphorylated EGFR (pEGFR, red). Scale bar: 50 μm.

A key finding was that *A. fumigatus* infection was predicted to activate the EGFR signaling pathway, altering the expression of 25 genes within this pathway (Fig. 1B). To verify that *A. fumigatus* activates EGFR in vivo, we used indirect immunofluorescence with a phosphospecific anti-EGFR antibody to detect the phosphorylation of EGFR in the lungs of mice with and without *A. fumigatus* infection. In uninfected mice, minimal phosphorylation of EGFR was detected in the pulmonary epithelial cells, whereas in mice infected with *A. fumigatus* there was substantial EGFR phosphorylation at foci of infection (Fig. 1C and S1A, B). When mice were treated with the EGFR inhibitor gefitinib, EGFR phosphorylation was decreased. Taken together, these results indicate that *A. fumigatus* infection activates EGFR signaling in the lung.

### EGFR mediates the response of small airway epithelial cells to *A. fumigatus in vitro*

Next, we investigated the interactions of *A. fumigatus* with EGFR *in vitro*, using the A549 cell line that resembles type II alveolar epithelial cells and the Tert-immortalized HSAEC1-KT human small airway epithelial (HSAE) cell line. By indirect immunofluorescence, we observed that EGFR accumulated around *A. fumigatus* hyphae in infected A549 and HSAE cells (Fig. 2A and S1C). Also, hyphae bound to EGFR in membrane protein extracts of A549 and HSAE cells (Fig. 2B). Although *A. fumigatus* infection inhibited the phosphorylation of EGFR (Y1068) in A549 cells, it stimulated EGFR phosphorylation in HSAE cells (Fig. 2C and D). The inhibition of EGFR phosphorylation in A549 cells was specific to *A. fumigatus* because incubating these cells with epidermal growth factor, a natural ligand of EGFR, stimulated strong EGFR phosphorylation (Fig. S1D).

**FIG 2.**
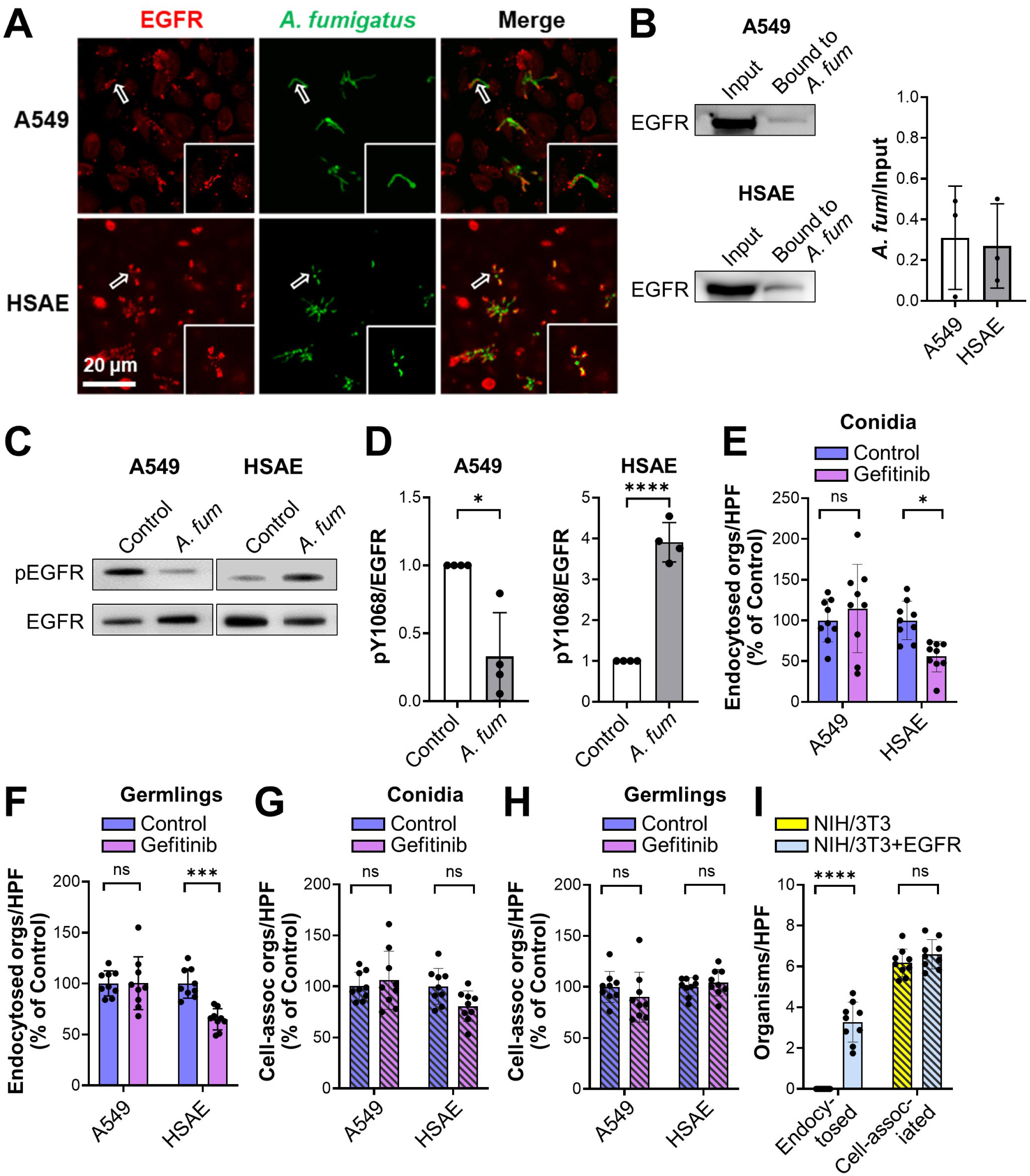
EGFR interacts with *A. fumigatus* and regulates fungal endocytosis by human small airway epithelial (HSAE) cells. (**A**) Confocal micrographs showing EGFR accumulation around *A. fumigatus* in A549 and HSAE cells after 2.5 h of infection. Hollow arrows indicate the organisms in the magnified images in the lower right of each panel. Scale bar 20 μm. (**B**) *A. fumigatus* binds to EGFR in membrane protein extracts of both A549 and HSAE cells. Representative immunoblots (right). Densitometric analysis of 3 immunoblots (left). Results are mean ± SD. (**C**) Western blots showing that *A. fumigatus* infection for 2.5 h inhibits EGFR phosphorylation in A549 cells but stimulates EGFR phosphorylation in HSAE cells. (**D**) Densitometric analysis of 4 phospho-EGFR Western blots such as the ones in (**C**). (**E** and **F**) Effects of the EGFR inhibitor, gefitinib on the endocytosis of *A. fumigatus* conidia (**E**) and germlings (**F**) by A549 and HSAE cells. (**G** and **H**) Effects of gefitinib on the cell-association (a measure of adherence) of *A. fumigatus* conidia (**G**) and germlings (**H**) with A549 and HSAE cells. (**I**) Endocytosis and cell-association of *A. fumigatus* by wild-type NIH/3T3 cells or NIH/3T3 cells that expressed human EGFR. Results in (**E-I**) are mean ± SD of 3 independent experiments, each performed in triplicate. *A. fum, A. fumigatus;* cell-assoc, cell-associated; orgs/HPF, organisms per high powered field; ns, not significant; \**P* < 0.05; \*\*\**P* < 0.001; \*\*\*\**P* < 0.0001 by unpaired Students’ t-test (**D**) or one way ANOVA with Dunnett’s test for multiple comparisons (**F-I**).

The functional significance of EGFR activation by *A. fumigatus* was assessed using a differential fluorescence endocytosis and cell association assay. Inhibition of EGFR with either gefitinib or an anti-EGFR antibody had no effect on the endocytosis of *A. fumigatus* conidia and germlings by A549 cells, whereas these treatments significantly reduced endocytosis of both morphotypes by HSAE cells (Fig. 2E and F, S1DEand F). EGFR inhibition had minimal effect on the number of conidia or germlings that were cell-associated with A549 or HSAE cells, a measure of adherence (Fig. 2 G and H, S1G and H).

To verify the role of EGFR in mediating the endocytosis of *A. fumigatus in vitro*, we tested the interactions of the fungus with the NIH/3T3 mouse fibroblast cell line. Germlings were poorly endocytosed by control NIH/3T3 cells that did not express EGFR but were efficiently endocytosed by NIH/3T3 cells that heterologously expressed human EGFR (Fig. 2I). The expression of EGFR on the NIH/3T3 cells did not significantly alter the adherence of *A. fumigatus* to these cells. Collectively, these results indicate that *A. fumigatus* activates EGFR both during pulmonary infection in immunosuppressed mice and during infection of HSAE cells, but not A549 cells. Also, activation of EGFR triggers the endocytosis of *A. fumigatus* by HSAE cells *in vitro*. Based on our findings that EGFR signaling altered the interactions of *A. fumigatus* with HSAE cells, but not A549 cells, we focused the remainder of our experiments on HSAE cells.

*A. fumigatus* both damages pulmonary epithelial cells and stimulates them to secrete proinflammatory cytokines and chemokines (26). We found that while inhibition of EGFR with gefitinib had no effect on fungal damage to A549 cells, it caused a slight, but statistically significant increase in damage to HSAE cells (Fig. S1I). Next, we assessed the effects of EGFR inhibition on the production of IL-1α, IL-1β, CXCL1, CXCL8, G-CSF, and GM-CSF uninfected and infected A549 and HSAE cells, using the same inoculum and time point for both cells. Under these conditions, A549 cells did not produce detectable amounts of IL-1α, IL-1β and G-CSF. *A. fumigatus* infection reduced the section of CXCL1, had no effect on the secretion of CXCL8, and stimulated the secretion of GM-CS F (Fig. S2). Although gefitinib reduced the basal secretion of CXCL1, it had no effect on the amount of CXCL1, CXCL8, GM-CSF induced by *A. fumigatus* infection. In contrast, *A. fumigatus* stimulated HSAE cells to secrete IL-1α, IL-1β, CXCL1, CXCL8, G-CSF, and GM-CSF, and both gefitinib and an anti-EGFR antibody inhibited the secretion of these inflammatory mediators (Fig. 3 and S3). Thus, EGFR signaling is required for *A. fumigatus* to induce a pro-inflammatory response in small airway epithelial cells *in vitro*.

**FIG 3.**
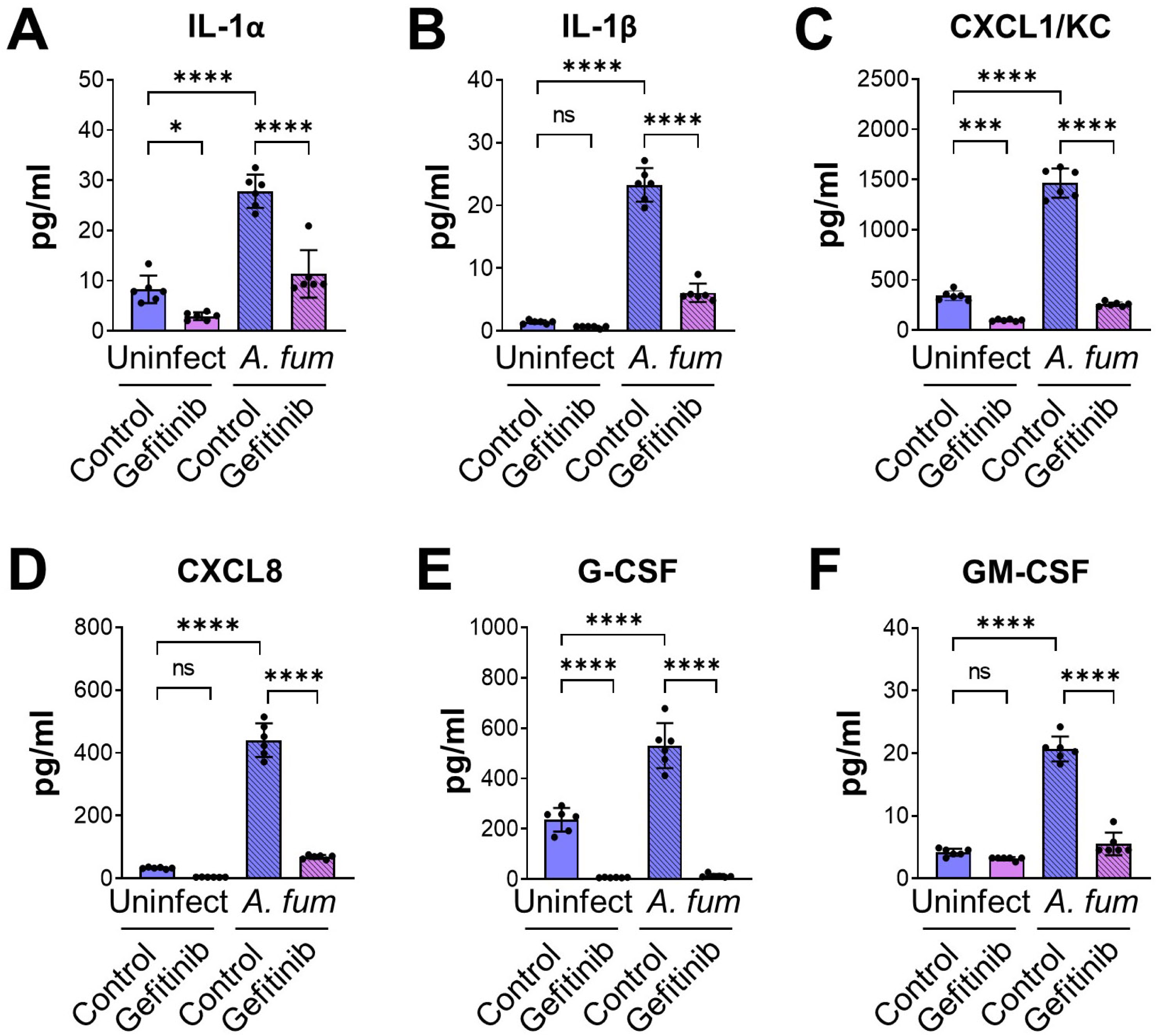
EGFR is required for maximal cytokine release in HSAE cells infected with *A. fumigatus*. (**A-F**) HSAE cells were treated with gefitinib or DMSO control, infected with *A. fumigatus* for 16 h, and then levels of the indicated cytokines were measured. Results are mean ± SD of 3 independent experiments, each performed in duplicate. *A. fum, A. fumigatus;* uninfect, uninfected; ns, not significant; \**P* < 0.05; \*\*\**P* < 0.001; \*\*\*\**P* < 0.0001 by one way ANOVA with the Dunnett’s test for multiple comparisons.

### EGFR is required for the host defense against *A. fumigatus* infection in immunosuppressed mice

In corticosteroid-treated mice, *A. fumigatus* infection induced EGFR phosphorylation in pulmonary epithelial cells, and this phosphorylation was inhibited by treating the mice with gefitinib (Fig. 1C). Treatment of the immunosuppressed mice with gefitinib prior to and during infection resulted in accelerated mortality and increased pulmonary fungal burden (Fig. 4A and B). In immunocompetent mice infected intratracheally with 10^8^ conidia, there was no mortality in either the control or gefitinib-treated mice (Fig. S3). Thus, the effect of EGFR inhibition on the outcome of invasive aspergillosis is dependent on the immune status of the host.

**FIG 4.**
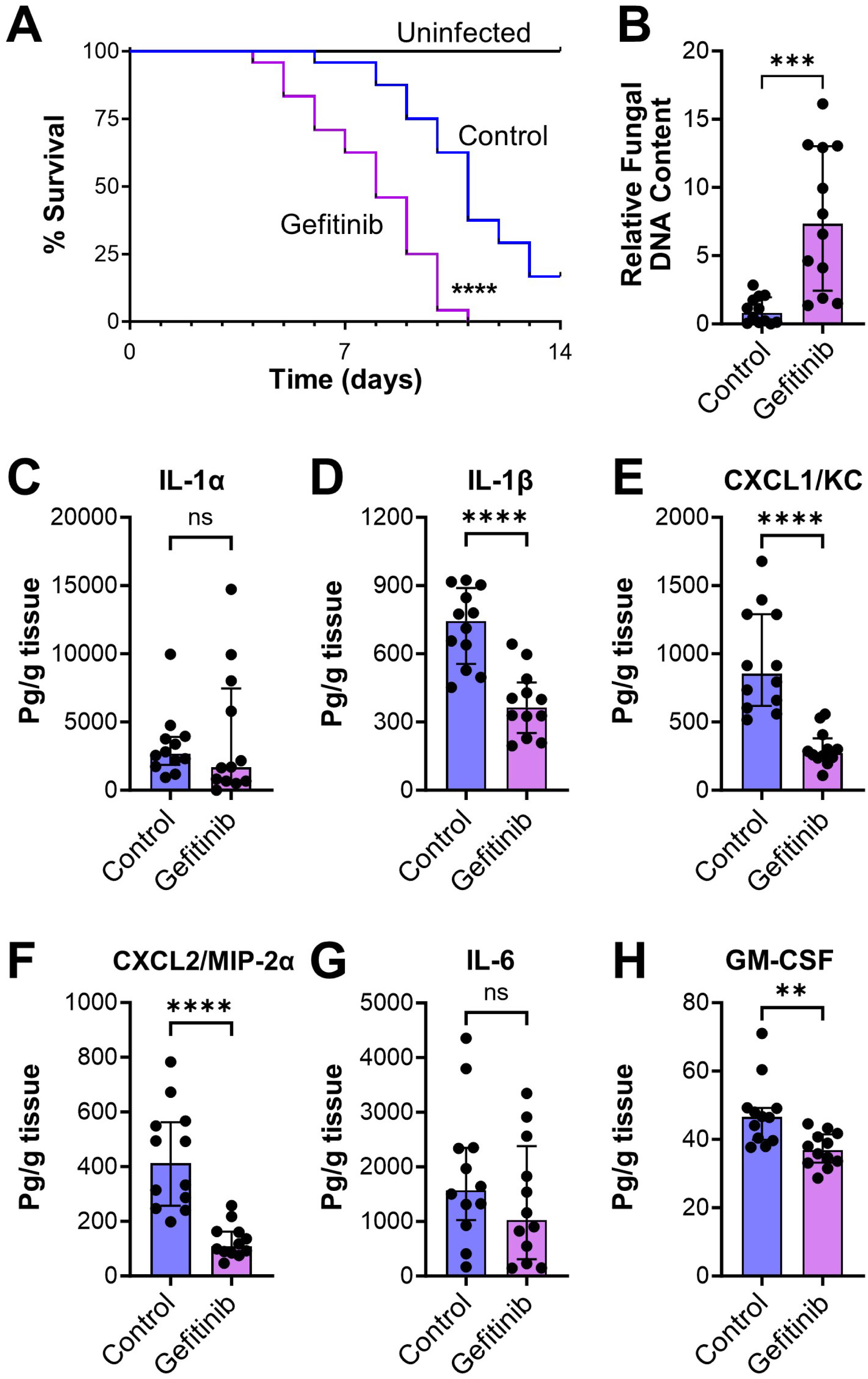
Gefitinib worsens outcome in immunosuppressed mice infected with *A. fumigatus.* (**A**) Effects of gefitinib on the survival of mice infected with *A. fumigatus* by aerosol inhalation. Results are the combined data from 2 experiments (n=24 mice per group). (**B**) Effects of gefitinib on pulmonary fungal burden after 4 d of infection, measured by the relative fungal DNA content in the infected mouse lungs. (**C-H**) Effects of gefitinib on the levels of the indicated inflammatory mediators in homogenates of the lungs of mice after 4 d of infection. Results in (**B-H**) are median ± interquartile range of 12 mice per group in a single experiment. ns, not significant, \*\**P* < 0.01, \*\*\**P* < 0.001, \*\*\*\**P* < 0.0001 by the log rank test (**A**) or Mann-Whitney test (**B-H**).

The deleterious effects of gefitinib in the immunosuppressed mice were likely due to impairment of the host inflammatory response because inhibition of EGFR reduced the production of IL-1β, CXCL1, CXCL2, and GM-CSF in the infected lungs (Fig. 4C-H). To analyze the effects of gefitinib on the phagocytosis and killing of *A. fumigatus* by phagocytes in the mouse lung, we used the FLARE technique (27, 28), in which mice were infected with *A. fumigatus* conidia that expressed dsRed and were labeled with AlexaFluor 633. In this assay, conidia that are alive exhibit fluorescence in both the dsRed and AlexaFluor 633 channels, whereas those that are killed no longer have dsRed fluorescence and only fluoresce in the AlexaFluor 633 channel.

We determined that although treatment of mice with gefitinib had no effect on the number of macrophages in the infected lungs, it caused a small, but significant reduction in *A. fumigatus* killing by these cells (Fig. 5A and S5). More importantly, gefitinib significantly reduced the accumulation of neutrophils and dendritic cells in the lungs but had no effect on the phagocytosis and killing of the organism by these cells (Fig. 5B and C). These results suggest that inhibition of EGFR reduces the production of proinflammatory mediators by pulmonary epithelial cells, which impairs the accumulation of neutrophils and dendritic cells into the lung and leads to decreased fungal killing.

**FIG 5.**
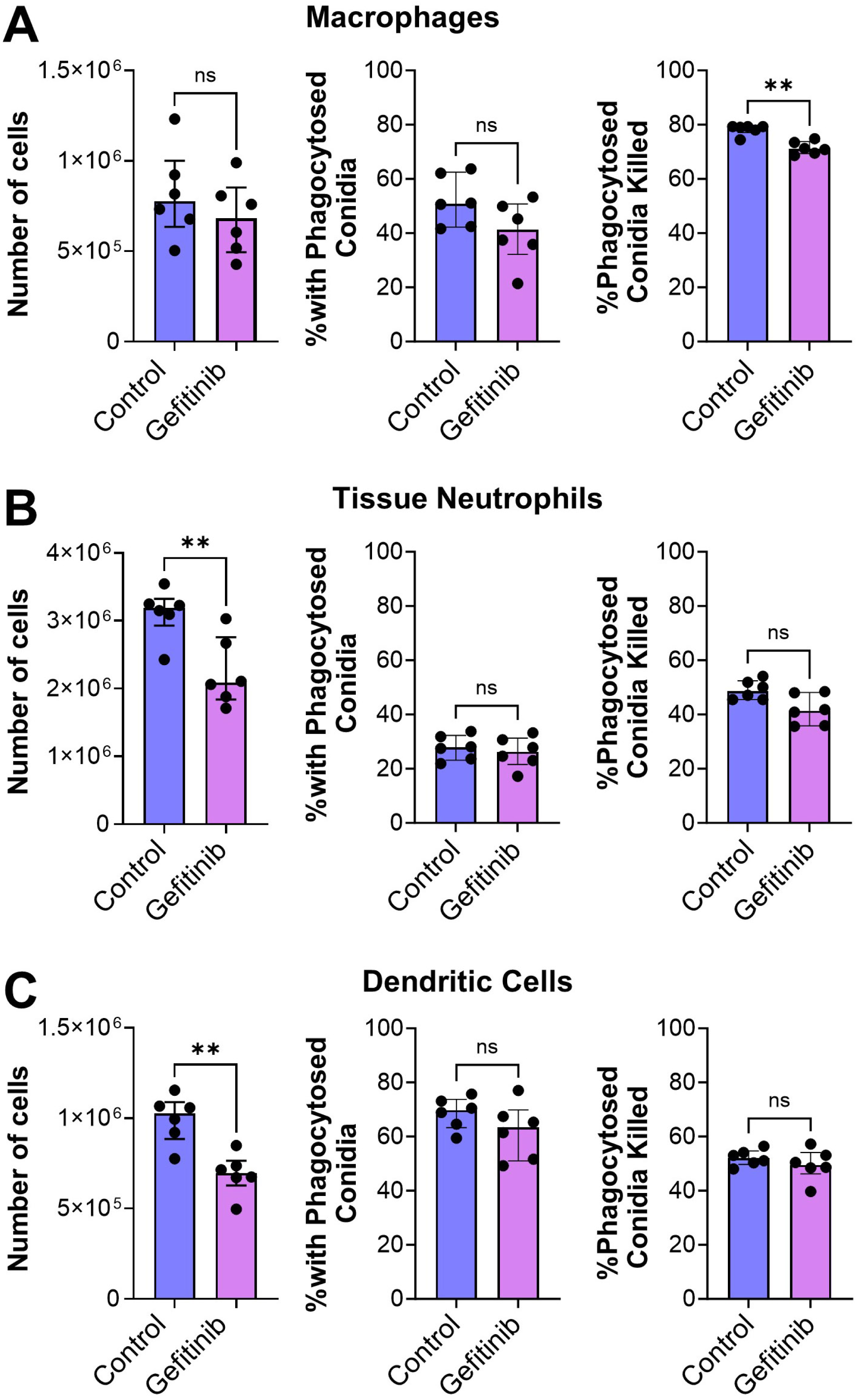
Gefitinib inhibits *A. fumigatus* killing by immune cells during invasive aspergillosis. (**A-C**) Corticosteroid-immunosuppressed mice were infected intratracheally with *A. fumigatus* conidia expressing DsRed and labeled with AlexaFluor 633. After 12 h, the tissue macrophages (**A)** neutrophils (**B**) and dendritic cells (**C**) were analyzed by flow cytometry to determine the number of cells in the samples, the percentage of cells with phagocytosed conidia, and the percentage of phagocytosed conidia that had been killed. Results are the median ± interquartile range of 6 mice per group in a single experiment. ns, not significant, \**P* < 0.05, \*\**P* < 0.01 by the Mann-Whitney test.

To test this model, we examined the effects of gefitinib on the chemotaxis of human neutrophils induced by medium conditioned by exposure to HSAE cells and/or *A. fumigatus.* Uninfected HSAE stimulated modest neutrophil chemotaxis, which was inhibited by treatment of the HSAE cells with gefitinib (Fig. 6). HSAE cells infected with *A. fumigatus* stimulated even more chemotaxis, which was reduced to basal levels by gefitinib. HSAE cells that has been incubated with killed *A. fumigatus* stimulated similar chemotaxis as was induced by uninfected HSAE cells. Thus, fungal viability is required for HSAE cells to stimulate maximal chemotaxis. Live, but not killed *A. fumigatus* grown in the absence of HSAE cells also induced strong chemotaxis. Importantly, this chemotaxis was not inhibited by gefitinib Collectively, these data indicate that gefitinib blocks the production of neutrophil chemotactic factors produced by HSAE cells. The finding that gefitinib had no effect on chemotaxis induced by live *A. fumigatus* alone also indicates that gefitinib has no direct effect on neutrophil chemotaxis.

**FIG 6.**
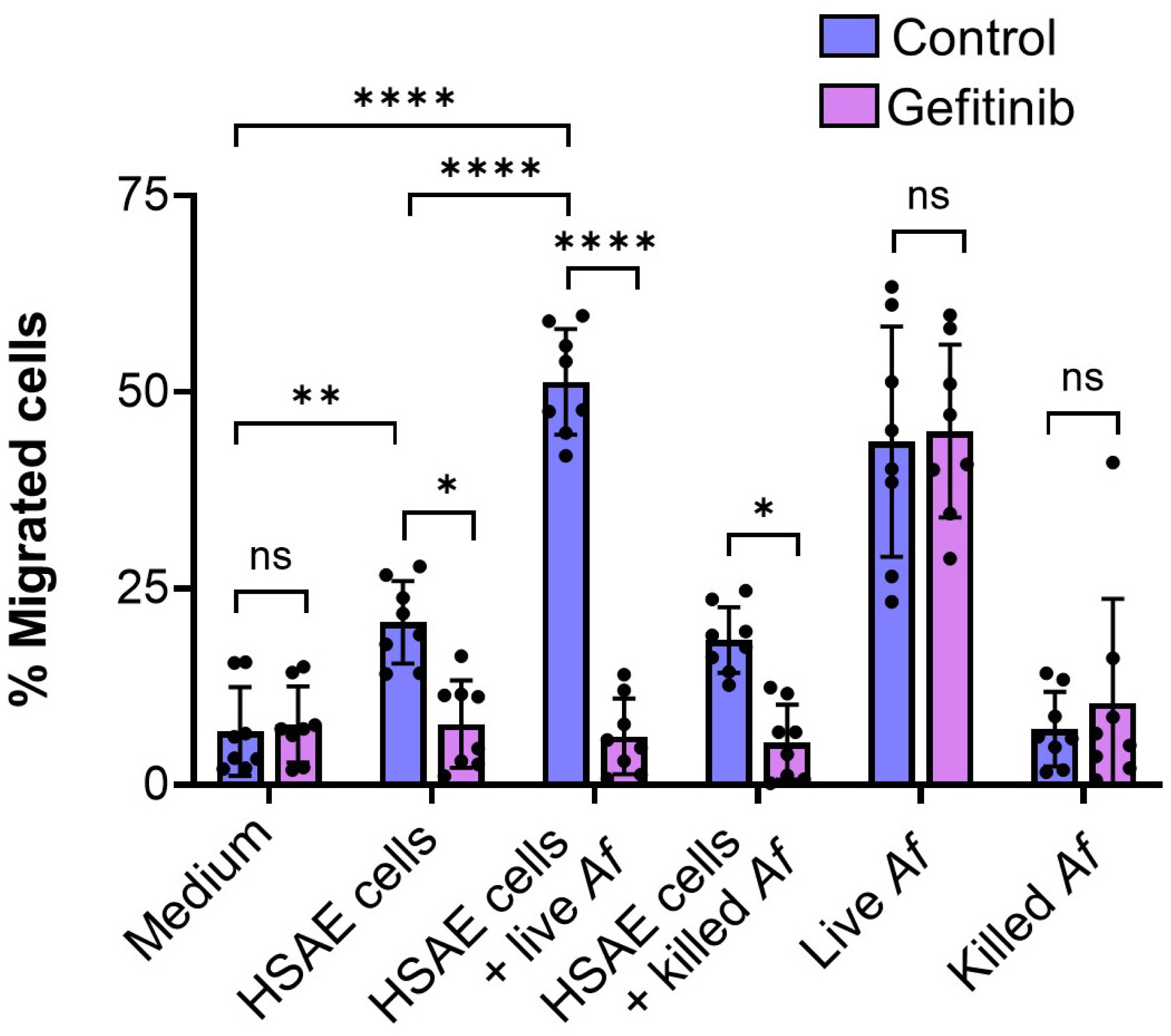
Gefitinib treatment of HSAE cells infected with *A. fumigatus* inhibits human neutrophil chemotaxis. Effects of gefitinib on neutrophil migration across transwell inserts induced by medium alone, and medium conditioned by uninfected HSAE cells, HSAE cells infected with live *A. fumigatus* (*Af*), HSAE cells incubated with paraformaldehyde killed *A. fumigatus*, live *A. fumigatus* or paraformaldehyde killed *A. fumigatus.* Results are the mean ± SD of 4 experiments each performed in duplicate. ns, not significant, \**P* < 0.05, \*\**P* < 0.01, \*\*\*\**P* < 0.0001 by one-way ANOVA with Sidak’s multiple comparisons test.

Next, we examined whether gefitinib affected the fungicidal activity of bone marrow-derived macrophages (BMDMs) *in vitro*. Although macrophages are known to express EGFR (29), exposure of these cells to gefitinib did not significantly alter their capacity to kill *A. fumigatus* after 8 and 16 h of infection (Fig. S6). Thus, it is likely that the modest reduction in the fungicidal activity of pulmonary macrophages caused by gefitinib *in vivo* was due to an indirect effect of the inhibitor, such as the reduction of proinflammatory cytokine production by the infected stromal cells.

## Discussion

It has been reported that some patients with cancer who are treated with EGFR inhibitors develop invasive aspergillosis (12–15). Our findings provide a potential explanation for these data. We determined that *A. fumigatus* activates EGFR signaling, both in HSAE cells *in vitro* and in the lungs of immunosuppressed mice with invasive aspergillosis. EGFR serves as a sensor for *A. fumigatus* in small airway epithelial cells. When the fungus interacts with these cells, EGFR is activated, leading to the production of proinflammatory cytokines and chemokines that recruit leukocytes and dendritic cells to foci of infection where they can kill the fungus.

It was notable that gefitinib treatment did not cause mortality in immunocompetent mice, even when they were infected with a very high inoculum of *A. fumigatus* Af293. It is possible that we might have observed accelerated mortality induced by gefitinib in mice infected with a strain of *A. fumigatus* such as CEA10 that is more virulent in mice (30). Nevertheless, it is likely that one or more EGFR-independent pathways can orchestrate the host inflammatory response in immunocompetent hosts. We speculate that the presence of these alternative pathways is the reason why invasive aspergillosis is not even more common in cancer patients treated with EGFR inhibitors. Our data indicate that when such alternative pathways are weakened or rendered inoperative by immunosuppression with high dose corticosteroids, the EGFR pathway plays a key role in generating the molecular signals that enhance the pulmonary defense against *A. fumigatus*.

We also found that the interaction of *A. fumigatus* with EGFR could trigger HSAE cells to endocytose the fungus *in vitro*. Previous studies have shown that epithelial cell invasion is a key step in the pathogenesis of invasive aspergillosis (4, 5). However, our results suggest that capacity of EGFR to trigger a proinflammatory response is more important for the pathogenesis of this disease than its ability to induce the endocytosis of this fungus.

These results are different from those with the Murcorales and *C. albicans*, where inhibition of EGFR with gefitinib protects mice from pulmonary mucormycosis and oropharyngeal candidiasis, respectively (9, 11). These previous studies demonstrated that EGFR is required for A549 cells to endocytose Mucorales fungi and for an oral epithelial cell line to endocytose *C. albicans.* Although the effects of gefitinib on the host inflammatory response to mucormycosis has not been investigated, it has been found that gefitinib significantly reduces the production of proinflammatory cytokines by oral epithelial cells infected with *C. albicans* (9). Taken together, these results suggest that EGFR functions as an epithelial receptor for multiple fungi that both triggers a proinflammatory response to and induces the endocytosis of fungal pathogens. During invasive pulmonary aspergillosis, blocking EGFR worsens disease by inhibiting the host inflammatory response, whereas during oropharyngeal candidiasis and pulmonary mucormycosis, blocking EGFR ameliorates disease by inhibiting epithelial cell endocytosis of the fungi.

A limitation of all studies of the role of EGFR in fungal infections in mice is that they have relied on small molecule inhibitors of EGFR such as gefitinib (9–11, 31), which might have off-target effects. The optimal approach to validate these inhibitor studies is to use mice in which EGFR has been genetically disrupted. It is known that global EGFR disruption in mice results in pre- or post-natal mortality, depending on the mouse strain (32, 33). However, it is probable that mice with conditional, site specific disruption of EGFR could be developed and used to test the role of this receptor in the pathogenesis of fungal infections.

At present, the mechanism by which *A. fumigatus* activates EGFR is unknown. It is has been determined that *C. albicans* stimulates EGFR via two mechanisms. *C. albicans* hyphae express the Als3 and Ssa1 invasins that interact with and directly activate EGFR (10). *C. albicans* hyphae also secrete the candidalysin pore forming toxin that activates EGFR indirectly by inducing the proteolytic cleavage and release of epiregulin and epigen, which are native EGFR ligands (31). Whether *A. fumigatus* activates EGFR by either of these mechanisms is a topic of active investigation.

## Materials and Methods

### Ethics statement

The mouse studies were approved by the Institutional Animal Care and Use Committee at the Lundquist Institute for Biomedical Innovation at Harbor-UCLA Medical Center. The mice were housed according to experimental group in HEPA-filtered laminar flow cages with unrestricted access to food and water. The vivarium is managed by the Lundquist Institute for Biomedical Innovation at Harbor-UCLA Medical Center in compliance with all polices and regulations of the Office of Laboratory Animal Welfare of the Public Health Service. The facility is fully accredited by the American Association for Laboratory Animal Care. The collection of blood from healthy volunteers was performed after obtaining informed consent. This protocol was approved by the Institutional Review Board of the Lundquist Institute for Biomedical Innovation at Harbor-UCLA Medical Center.

### Strains and growth condition

The *A. fumigatus* Af293 strain was purchased from the American Type Culture Collection (ATCC). The Af293-DsRed strain was a gift from Tobias M Hohl, Memorial Sloan Kettering Cancer Center (27). The Af293-GFP strain was constructed as previously described (4). All *A. fumigatus* strains were grown on Sabouraud dextrose agar (Difco) at 37°C for 7-10 days prior to use. Conidia were harvested by gently rinsing the agar plates with PBS containing 0.1% Tween 80 (Sigma-Aldrich), after which the conidia were filtered through a 40 µm cell strainer (Corning) and enumerated with a hemacytometer. To observe the host cell interactions with live germlings, the conidia were pre-germinated in Sabouraud dextrose broth (Difco) at 37°C for 5.5 h.

### Cell lines

A549 cells were purchased from the ATCC and cultured in F-12 K medium (ATCC, 30-2004) supplemented with 10% fetal bovine serum (FBS) (Gemini Bio-Products) and 1% streptomycin and penicillin (Irvine Scientific). HSAEC1-KT (HSAE) cells were obtained from the ATCC and cultured in SAGM BulletKit medium (Lonza CC-3119 and CC-4124). The NIH/3T3 cell lines with or without EGFR expression were provided by Nadege Gaborit (Institut de Recherche en Cancérologie de Montpellier, Montpellier, France) (34) and cultured in Dulbecco’s Modified Eagle Medium (DMEM) supplemented with 10% FBS and 1% streptomycin and penicillin. All host cells were cultured at 37 °C in 5% CO_2_. The A549 and HSAEC1-KT cell lines were authenticated by the ATCC, the NIH/3T3 wild-type and EGFR expressing cell lines were authenticated by immunoblotting to detect the absence and presence of EGFR, respectively. All cell lines had no mycoplasma contamination.

### Mouse models of invasive aspergillosis

To study the host transcriptional response during invasive aspergillosis, we used our standard corticosteroid immunosuppressed mouse model (35, 36). Briefly, 6-week male Balb/C mice (Taconic Laboratories) were immunosuppressed with cortisone acetate at 500 mg/kg administered subcutaneously every other day, starting on day -4 relative to infection and finishing on day +4. To prevent bacterial infections, enrofloxacin (Baytril, Western Medical Supply) was added to the drinking water at a final concentration of 0.005% on day -5 relative to infection. The mice were infected by placing them in a chamber into which 1.2×10^10^ conidia were aerosolized. Control mice were immunosuppressed but not infected. After 2, 4, and 6 days of infection, 3 mice were sacrificed. Their lungs were harvested and snap frozen for RNA extraction.

To assess the effects of gefitinib on the outcome of infection, the mice were immunosuppressed with cortisone acetate as above except that the dose was reduced to 250 mg/kg to prolong survival so that the deleterious effects of gefitinib were more apparent. The mice were fed powdered chow with or without 0.2 mg gefitinib per g beginning on day -3 before infection and continuing for the duration of the experiment. For the survival experiments, mice that were immunosuppressed and treated with gefitinib, but not infected were used as the uninfected control. The mice were monitored for survival twice daily by an unblinded observer and moribund mice were humanely euthanized. The survival experiments were repeated twice, and the results were combined for a total of 24 mice per group. The effect of gefitinib on the survival of immunocompetent mice was determined by inoculating 8 mice per group intratracheally with 10^8^ *A. fumigatus* conidia.

To determine the effects of gefitinib treatment on pulmonary fungal burden and cytokine levels, 12 mice per group were immunosuppressed, treated with gefitinib, and infected in the aerosol chamber. After 4 d of infection, the mice were sacrificed and their lungs were harvested and homogenized in lysis solution (ZR Fungal/Bacterial DNA MiniPrep^TM^, Epigenetics) using a gentleMACS dissociator (Miltenyi Biotec). The homogenates were clarified by centrifugation and the cytokine content of the supernatant was determined using a Luminex Multiplex Assay. The DNA in the pellet was isolated using the ZR Fungal/Bacterial DNA MiniPrep^TM^ kit following the manufacturer’s directions. The fungal DNA content in the various samples was quantified by real-time qPCR using primers ASF1 (5-GCACGTGAAATTGTTGAAAGG-3) and ADR1 (5-CAGGCTGGCCGCATTG-3) using the 2^-ΔΔCT^ method with mouse GAPDH (5-CAACAGCAACTCCCACTCTTC-3 and 5-GGTCCAGGGTTTCTTACTCCTT-3) as the reference gene.

To detect EGFR phosphorylation *in vivo*, mice were immunosuppressed with cortisone acetate (250 mg/kg) and treated with gefitinib as above. They were anesthetized with xylazine and ketamine and inoculated by intratracheal injection with 3 x 10^7^ conidia of *A. fumigatus* Af293. After 12 h of infection, 3 mice from each group were sacrificed and the lungs were perfused by injecting 5 ml of PBS into the right ventricle. Next, the lung were harvested and processed for immunohistology as described below. In similar experiments, the FLARE technique was used to determine *A. fumigatus* phagocytosis and killing by various leukocyte subsets (27). The mice were immunosuppressed, treated with gefitinib, and infected as above except that there were 6 mice per experimental group and the mice were inoculated with 3 x 10^7^ dsRed expressing conidia that had been labeled with AlexaFluor 633. After 12 h of infection, the mice were sacrificed and their lungs were used for the FLARE assay as described below.

### RNA-seq analysis

To extract RNA from the mouse lungs, each lung sample was placed in a lysing matrix C tube (6912050, MP Biomedicals) containing a single 0.25 inch diameter ceramic sphere (6540424, MP Biomedicals) and homogenized with a bead beater (FastPrep FP120, Thermo Scientific). The RNA was isolated using the RiboPure RNA kit (AM1924, Invitrogen) following the manufacturer’s instructions and contaminating DNA was removed using the Turbo DNA-free kit (AM1907, Invitrogen).RNA-seq libraries were prepared using the TruSeq RNA Sample Prep kit (Illumina, San Diego, CA) per the manufacturer’s protocol. Next, 100 bp paired end reads were generated using the Novaseq platform (Illumina, San Diego, CA). From each of the 18 sequencing libraries, we obtained an average of 101.2 ± 15.9 million reads that were mapped to the mouse reference genome using HISAT2, a short read aligner (Table S5). The work presented here focuses our analysis of the host transcriptome response. The *A. fumigatus* response will be presented elsewhere. A gene was considered differentially expressed if the absolute fold-change was ≥ 2.0 and the false discovery rate was ≤ 0.05. To identify host signaling pathways that were activated or inhibited by *A. fumigatus* infection, the Upstream Regulator Analytic from the Ingenuity Pathway Analysis software (Ingenuity Systems) was used as previously described (37).

All of the raw sequencing reads from this study are available at the NCBI Sequence Read Archive (SRA) under BioProject accession number PRJNA1140223. The specific sample accession numbers are presented in Table S4.

### Indirect immunofluorescence

To detect the effects of *A. fumigatus* infection on the phosphorylation of EGFR *in vivo*, the mouse lungs were snap frozen in Optimal Cutting Temperature (OCT) tissue embedding medium (4585, Fisher Scientific), after which 9-µm thick sections were cut with a cryostat, dried for 1 h, and fixed with ice cold methanol. The cryosections were rinsed with PBS, blocked with 1% goat serum, and then strained with AlexaFluor 568-labeled anti-EGFR-p-Tyr1068 (2234, Cell Signaling), AlexaFluor 488-labeled anti-*Aspergillus,* and BV421-conjugated anti-CD326 (563214, BD Biosciences) antibodies. Control sections were stained with AlexaFluor 568-labeled rabbit IgG instead of the anti-EGFR antibody. The cells were imaged by confocal microscopy (Leica SP8) and the same image acquisition settings were used to enable comparison of fluorescence intensity among the different samples.

The accumulation of EGFR around *A. fumigatus* hyphae during infection *in vitro* was imaged as our previously described methods (4). A549 cells or HSAE cells were infected with 1×10^5^ germlings of GFP expressing *A. fumigatus* Af293 for 2.5 h. Cells were fixed with 4 % paraformaldehyde, permeabilized with 0.1% triton X100, blocked with 5% goat serum (vol/vol), and stained with an anti-EGFR antibody (4267, Cell Signaling) followed by AlexaFluor 568-labeled goat anti-rabbit antibody (A11036, Invitrogen). The cells were imaged as described above.

### FLARE assay

After disrupting the excised lungs with a gentleMACs tissue dissociator (Miltenyi Biotec) to yield single cells, the red blood cells were removed by treatment with ACK lysis buffer (Thermo Fisher). The resulting single-cell suspension was counted and 2×10^6^ cells were used for antibody staining. The antibodies used were BUV395-anti-CD45 (564279, BD), FITC-anti-Ly6G (11-9668-82, Invitrogen), Pe-Cy7-anti-Ly6C (560593, BD), FITC-anti-CD103- (11-1031-82, Invitrogen), Pacific Blue-anti-CD11b (NC9908052, Biolegend), Ly6B.2-AF700 (NBP213077Z), CD11C-PE-Cy7 (558079, BD), MHCII-AF700 (56-5321-82, Invitrogen). The staining protocols were followed as previously described. The neutrophils were identified as CD45^+^Ly6G^+^Ly6C^lo^CD11b^+^Ly6B.2^+^, macrophages as CD45^+^CD11c^+^MHCII^var^ and dendritic cells as CD45^+^CD11c^+^MHCII^var^CD103^-^CD11b^+^. Data were acquired using a five-laser BD FACSymphony™ A5 and analyzed with Flowjo Software V10. The gating strategy is diagrammed in Fig. S4.

### EGFR pull down assay

The capacity of *A. fumigatus* cells to affinity purify EGFR from lysates of A549 and HSAE cells was determined as previously described (4). Briefly, 2×10^7^ *A. fumigatus* germlings were incubated with 2.5 µg total epithelial cell proteins in 1.5% octyl-glucopyranoside in PBS for 1 h on ice. After extensive rinsing, the proteins that remained bound to the germlings were eluted with 6 M urea and separated by SDS-PAGE. The presence of EGFR in these proteins was detected by Western blotting with an anti-EGFR antibody (4267, Cell signaling).

### Western blotting detection of EGFR phosphorylation *in vitro*

Host cells were cultured in 24-well tissue culture plates and infected with 1×10^6^ germlings for 2.5 h. For EGF stimulation, the cells were incubated with 20 ng/ml EGF for 5 min. Next, the cells were rinsed with HBSS containing protease and phosphatase inhibitor cocktails, and then lysed with sample buffer. The lysates were collected and boiled in an equal volume of sample buffer. The proteins in the lysates were separated by SDS-PAGE, and phosphorylated EGFR was detected by Western blotting with a phospho-specific anti-EGFR antibody (2234, Cell Signaling). The blots were then stripped, and the total EGFR was detected with anti-EGFR antibody (4267, Cell Signaling). The experiments were repeated 4 times and densitometric analysis of the blots was performed using Image studio Lite 5.2 (Li-Cor).

### Host cell association and endocytosis assay

The cell association and endocytosis of *A. fumigatus* conidia and germlings by A549, HSAE or NIH/3T3 cell lines was determined by our previously described differential fluorescence assay (4). Host cells were grown to confluency on glass coverslips in 24-well tissue culture plates and then infected 1×10^5^ conidia or germlings of Af293-GFP per well. Conidia were incubated with host cells for 6 h and germlings for 2.5 h. After the incubation period, the cells were rinsed with 1 ml of PBS, fixed with 4% paraformaldehyde, and stained with a rabbit anti-*Aspergillus* polyclonal antibody (B65821R, Meridian Life Science, Inc.) followed by an AlexaFluor 568-labeled goat anti-rabbit antibody (A11036, Life Technologies). The coverslips were mounted inverted onto microscope slides and GFP (green) or AlexaFluor 568 (red) labeled organism were viewed using an epifluorescence microscope. At least 100 organisms per coverslip were scored and each strain was tested in triplicate in three independent experiments.

### Cell damage release assay

To quantify *A. fumigatus-*induced damage to HSAE cells, the cells were incubated with ^51^Cr overnight in 24-well tissue culture plates. Next day, the unincorporated ^51^Cr was removed by washing the cells twice with HBSS. The cells in each well were infected with 5 x 10^5^ conidia in 1 ml of SAGM BulletKit medium. After 20 h, the top 0.5 ml of the medium in each well was collected and the ^51^Cr content was measured by gamma counting. The remain medium was collected and cells were lysed with 6 N NaOH for 15 min followed by rinsing with RadiacWash (Biodex Medical Systems, Inc.) two times. The ^51^Cr content of the remaining medium, cell lysate and RadiacWash was also measured by gamma counting. Uninfected HSAE cells were processed in parallel to calculate the spontaneous release. The percent specific release of ^51^Cr was calculated as previously described (4). Each experiment was performed in triplicate and repeated three times.

### Cytokine and chemokine analysis

The cytokine levels in cell culture supernatants and lung homogenates were determined using the Luminex Multiplex Assay (R&D Systems). For *in vitro* studies, HSAE cells were grown to confluency in 24-well tissue culture plates. Each well of cells was infected with 5×10^5^ conidia of *A. fumigatus* Af293 in 1 ml of SAGM BulletKit medium. After 18 h of infection, 500 ml of the supernatant was collected, centrifugated, and stored in aliquots at -80 °C. At a later date, the cytokine content in the conditioned media were determined using the Luminex kit. The experiments were performed in duplicate and repeated three times.

For *in vivo* studies, the mouse lungs were harvested and homogenized in 1 ml of PBS containing 10 µl of proteinase inhibitor cocktail (P8340, Simga-Aldrich). The homogenates were centrifuged and stored at -80 °C for later cytokine analysis. The concentration of IL-1α, IL-1β, CXCL1, CXCL2, and IL-6 were determined using the Luminex assay. The concentration of GM-CSF was determined by ELISA (MGM00, R&D Systems).

### EGFR inhibition studies

To determine the effect of EGFR inhibitors on epithelial cell endocytosis, damage, and cytokine production, the host cells were incubated with 0.1 µM gefitinib (S1025, Selleck Chemicals) in 0.1% DMSO. Control cells were incubated with 0.1% DMSO in parallel. For antibody blocking experiments, the host cells were incubated with 10 µg/mL of cetuximab (Erbitux, ImClone Systems) and a similar concentration of mouse IgG was used as a control. Both gefitinib and the anti-EGFR antibody were added to the cells 45 min prior to infection with *A. fumigatus* and remained in the medium for the duration of the experiment.

### Neutrophil Chemotaxis assay

To test neutrophil chemotaxis towards media conditioned by *A. fumigatus* and HSAE cells, blood was collected from healthy volunteers by venipuncture and mixed with K_3_EDTA (#E-0270, Sigma-Aldrich). The neutrophils were isolated using gradient separation with Histopaque 1077 (#10771, Sigma-Aldrich) and Histopaque 1119 (#11191, Sigma-Aldrich) following the manufacturer’s instructions. The cells were washed once with HBSS without Ca^++^ or Mg^++^ (21-022-CV, Corning), suspended in RPMI 1640 medium with L-glutamine (9161, Irvine Scientific) containing 10% pooled human serum (#100–110, Gemini Bioproducts, Inc.) and enumerated using a hemacytometer. To test the effects of gefitinib on chemotaxis, the neutrophils were incubated with 0.1 μM gefitinib for 20 min prior to performing the assay. The gefitinib remained in the medium for the duration of the experiment. Control cells were incubated in 0.1% DMSO in parallel.

For the cell migration assay, 2×10^5^ neutrophils were seeded onto 24-well inserts containing a polycarbonate membrane with 5.0 μm pores (3421, Corning) that had been coated with 10 mg/ml fibronectin (33016015, Gibco). The neutrophils were allowed to settle for 20 min, after which the inserts were placed into a 24-well tissue culture plate containing the chemotactic stimuli. For chemotaxis stimulated by *A. fumigatus* alone, 5 x10^5^ germlings in SAGM BulletKit medium were added to the 24-well plate and incubated for 2.5 h, after which the filter inserts containing neutrophils were added to these wells. For chemotaxis stimulated by infected HSAE cells, the cells were infected with *A. fumigatus* in the presence and absence of gefitinib for 18 h as in the cytokine experiments. The conditioned medium was collected, filter sterilized, and stored at -80°C for later use. On the day of the experiment, the conditioned medium was warmed to 37°C and added to the 24-well plate immediately before adding the filter inserts. After a1-h incubation, the number of neutrophils that had migrated to the bottom of the wells was quantified by using the CellTiter-Glo assay (G7570, Promega) following the manufacturer’s instructions. Wells containing *A. fumigatus* in the absence of PMNs were processed in parallel to control for the minimal ATP release by the fungal cells. The experiment was performed in duplicate and repeated 4 times with separate neutrophil donors.

### BMDM killing assay

The effects of gefitinib on the fungicidal activity of macrophages were determined using a colony counting assay. BMDMs were isolated from the femurs of 6-week-old Balb/c mice (Taconic Laboratories) and differentiated by growth in (BioLegend) in Dulbecco’s Modified Eagle’s Medium (DMEM) (ATCC) with 10% fetal bovine serum (Gemini Bio-Products), and 1% streptomycin and penicillin, supplemented with 150 ng/ml M-CSF. After 7 days, the differentiated cells were harvested and 2 x 10^5^ cells were added to 24-well tissue culture plates. The next day, cells were treated with gefitinib or 0.1% DMSO alone for 45 min and 1 x 10^4^ *A. fumigatus* conidia was added. After 8 h and 18 h incubation, the BMDMs were lysed with sterile distilled water. The lysates were collected and sonicated and then the number of viable organisms was determined by quantitative culture. A similar number of conidia was inoculated without BMDMs as the initial inoculum control. Each experiment was performed in triplicate and repeated three times.

### Statistics

All statistical analyses were performed in GraphPad Prism version 10. The *in vitro* data were analyzed using the unpaired Student *t* test or one-way ANOVA with Dunnett’s test for multiple comparisons and presented as mean ± SD. The *in vivo* data were analyzed with the Mann-Whitney test and presented as median ± interquartile range. *P* values ≤ 0.05 were considered as statistically significant.

## Acknowledgements

This work was supported by NIH grants R01AI162802 (SGF), U19AI110820 (VMB, SGF), and R01AI141360 (VMB).

---

**Supplementary Table 1.** Differentially expressed host genes after 2 days of infection.

**Supplementary Table 2.** Differentially expressed host genes after 4 day of infection.

**Supplementary Table 3.** Differentially expressed host genes after 6 day of infection.

**Supplementary Table 4.** Upstream Regulator analysis of host pathways in response to *A. fumigatus*.

**Supplementary Table 5.** RNA-seq mapping statistics for each sample.

